# Aurora vent field is a hotspot for microbial hydrogen oxidation in the Arctic Ocean

**DOI:** 10.64898/2026.02.17.706317

**Authors:** Emily Olesin Denny, Petra Hribovšek, Samuel I. Pereira, Claudio Argentino, Giuliana Panieri, Achim Mall, Francesca Vulcano, Runar Stokke, Eoghan P. Reeves, Ida H. Steen, Håkon Dahle

**Author notes:** Corresponding author: Emily Olesin Denny.

## Abstract

Molecular hydrogen (H_2_) is a widespread, energetically efficient reductant supporting microbial metabolism across most known ecosystems. Although seafloor hydrothermal vents are major energy providers for H_2_-oxidizing microorganisms, the diversity of H_2_ oxidation potential in H_2_-rich systems remains poorly constrained. Here, we use a metagenomic approach to, for the first time, assess the genome-resolved microbial energy conservation potential within hydrothermal deposits and sediments from the ice-covered, extraordinarily H_2_-rich Aurora Vent Field in the Arctic Ocean. Community-wide analysis revealed broad taxonomic representation of microorganisms with the potential to consume H₂ for energy conservation. Notably, we report the first genome belonging to the cosmopolitan Zetaproteobacteria genus *Mariprofundus* encoding the capacity for H_2_ oxidation. Additionally, novel, highly abundant Aquificota at Aurora encode uptake hydrogenases not previously characterized as central to H_2_ oxidation at deep sea vents. The encoded gene content of abundant taxa points to a preference for flexible rather than obligate lithotrophic energy metabolism. A substantial fraction of inferred H₂-oxidizing potential is associated with presumed heterotrophs, potentially enhancing carbon transfer efficiency within the Aurora microbial food web. Overall, this study sheds new light on the importance of H_2_ availability for shaping microbial communities in hydrothermal systems.

**Importance:** Hydrogen oxidation is a potent energy source for microorganisms that contributes to shaping and sustaining food webs in marine and terrestrial habitats, with particular importance in deep-sea chemosynthetic communities. This study provides detailed insights into how exceptionally high hydrogen availability at the Aurora Vent Field may influence microbial community structure, revealing that hydrogen oxidation potential is widespread among diverse, metabolically versatile taxa rather than obligate hydrogen oxidizers. Our findings refine current understanding of microbial energy flow and carbon cycling in hydrogen-rich hydrothermal systems. These results inform future efforts to model microbial food webs, predict ecosystem responses to changing geochemical conditions, and explore metabolic interactions in deep-sea chemosynthetic habitats more broadly.

## Introduction

Deep-sea hydrothermal vents are ancient ecosystems sustained by chemolithotrophic primary production that is tightly coupled to availability of geochemically-derived reductants ^1^. At the seafloor, mixing of high-temperature, reduced hydrothermal fluids with oxygenated seawater generates strong chemical disequilibria that provide chemical energy for anabolic and catabolic processes in chemotrophic primary producers ^2,3^.

Molecular hydrogen (H_2_) in particular is widely present in hydrothermal systems and is a highly efficient and versatile reductant for energy acquisition via lithotrophic processes ^2,4–8^. However, most hydrothermal vents discovered to date worldwide appear influenced by alteration of mafic- and felsic igneous rocks in the subseafloor, and consequently the vast majority of the several hundreds of characterized hydrothermal vent fields emit fluids with low and variable H_2_ concentrations, typically well below ∼5 mM ^9,10^. H_2_ concentrations above this have, in contrast, only been reported in a handful of systems (< ∼10), primarily those affected by serpentinization of ultramafic lithologies (e.g. Charlou et al. (2010) ^11^). Even after extensive mixing of vent fluids with oxic seawater (e.g. dilution factors of 10^2^ - 10^4^) as occurs in rising and non-buoyant plumes ^9^, H_2_ availability in such hydrothermal systems remains orders of magnitude higher than in ambient seawater ^12–14^. Research in the last few decades has made it increasingly clear that oxidation of trace atmospheric H_2_ can be an important energy source for sustaining microbial communities in H_2_-poor habitats such as soils, where H_2_ oxidation serves as a supplemental energy source for organoheterotrophic microorganisms ^15^. Together, these observations highlight the potential for H_2_ oxidation to fuel not only primary producing lithoautotrophs in hydrothermal systems, but also microorganisms associated with organoheterotrophic consumption.

Microbial utilization of H_2_ as an energy source requires the production of uptake hydrogenases. Hydrogenases are broadly classified, based on metal centre composition, into Ni,Fe hydrogenases, Fe,Fe hydrogenases and Fe hydrogenases. Among these, the Ni,Fe hydrogenases include those responsible for H_2_ oxidation – the uptake hydrogenases. These hydrogenases belong to Ni,Fe subgroups 1a-1l and 2a, which vary in their electron acceptors, affinities for H_2_ and tolerance of oxygen ^15^. Ni,Fe uptake hydrogenases are widely distributed within bacteria and archaea that obligately, facultatively, or mixotrophically gain energy through H_2_ oxidation ^16^. Little is known about the distribution of hydrogenases (including uptake hydrogenases) in hydrothermal systems. Meta-omic studies conducted thus far indicate that low-affinity Ni,Fe uptake hydrogenases of subgroups 1b and 1d are the main acting H_2_ oxidizing catalysts encoded within vent microbial communities ^17–20^. However, systematic surveys of the breadth of uptake hydrogenases present in hydrothermal systems are still lacking, restricting our understanding of the full H_2_ oxidation potential among vent-associated microbial community members.

The perennially ice-covered Aurora Vent Field (AVF), located near the western end of the Gakkel Ridge in the Arctic Ocean, represents a unique marine habitat for investigating how extremely elevated H_2_ availability can shape microbial communities ^21–25^. The first visually guided (ROV-based) vent fluid and deposit sampling at AVF, however, was only conducted in October 2021 during the second HACON research expedition ^23,25^. Investigation of the geochemistry of high-temperature ‘black smoker’ fluids (350 – 356 °C) at AVF in 2021 revealed uniformly high H_2_ concentrations in endmember fluids that greatly exceed all other vent fluids measured to date, approaching 0.05 M ^25,26^. In 2023, follow-up exploration of other nearby black smoker vents within the field yielded a single vent fluid sample with a similarly high temperature and H_2_ abundance ^27^, suggesting that such extreme H_2_ contents in AVF source fluids are stable, at least over several years.

Here we present, using a metagenomic approach, the first characterization of the community structure and genomic potential of microbial assemblages found at the H_2_-rich AVF. Based on the relative endmember concentrations in the AVF fluid (47.5 mM H_2_ and *ca.* 1 mM H_2_S and CH_4_), H_2_ oxidation is highly likely the primary energy source sustaining life at AVF, followed by sulfide and methane oxidation ^25–27^ .Thermodynamic assessments of other ultramafic-influenced vent fields (Rainbow, Logachev, Kairei) with much lower endmember H_2_ (<16 mM), but comparable CH_4_ and H_2_S, clearly demonstrate H_2_ oxidation is a strongly dominant energy source for primary producers when vent fluid extensively mixes with seawater ^2,21^.

Although this work focuses on H_2_ oxidation potential at AVF, we also elucidate metagenomic signatures of other geochemical cycling to provide a holistic report of the functional potential within microbiological communities of this newly described and highly inaccessible hydrothermal vent field. Overall, this extreme ecosystem appears to be a promising environment for gaining new insights into microbial communities driven by H_2_ oxidation and for encountering novel H_2_ oxidizers.

## Results

### Sampling and DNA extraction

ROV exploration of AVF during the HACON21 expedition revealed many hydrothermally-impacted surfaces with iron oxide-like or sulfide-related discoloration near the three main vigorously venting black smoker structures. These structures have been described extensively elsewhere ^23,25^. During five ROV dives a total of 70 samples of chimney wall precipitates, hydrothermal deposits, and sediment cores were taken near these sites of active venting (Fig. 1, Supplementary Table S1).

**Fig. 1.**
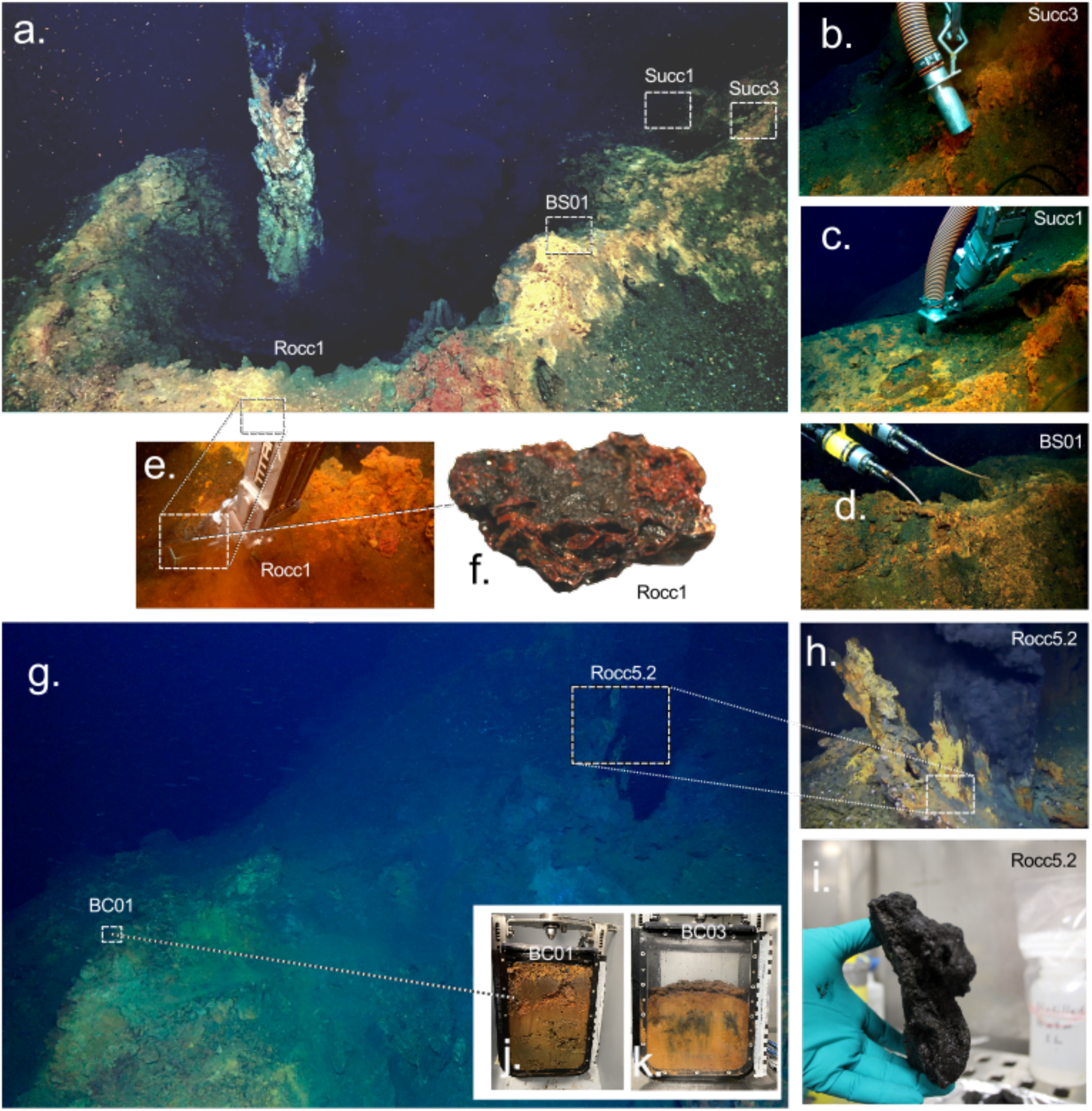
Overview of sampling locations at the Aurora Vent Field (AVF) where DNA was detected. **(a)** Visual of all sampling sites on the sulfide mound surrounding Hans Tore vent. **(b–c)** Still images of suction sampling (Succ1 and Succ3). **(d)** Hydraulic cylinder sampling (BS01). **(e–f)** Hydrothermal deposit collection (Rocc1) and the corresponding sample photographed onboard the vessel. **(g)** Sampling areas near Enceladus vent. **(h–i)** Collection and onboard image of an inactive chimney hydrothermal deposit near the Enceladus black smoker (Rocc5.2). **(j–k)** Shallow sediment cores collected a few meters from the Enceladus vent (BC01 and BC03).

All attempted DNA extractions from samples of active black smoker sulfide chimney deposits yielded DNA concentrations below the detection limit and therefore were not processed further. This included samples representing only the outer few mm of active chimney wall material, where sulfides are exposed to seawater (Supplementary Table S1) – regions typically highly enriched in microbial biomass in hydrothermal chimneys elsewhere ^28–30^. Notably, other vent fields on the Arctic Mid-Ocean Ridges (AMOR), such as the Loki’s Castle, Fåvne, and Jan Mayen vent fields, exhibit cm-thick, clearly visible biofilms that are widespread on vent structures ^20,31–33^. The lack of quantifiable DNA on AVF chimneys is supported by Lapointe et al. (2025) ^25^, who reported that these active chimney structures were unusually thin-walled and dominated by high-temperature minerals, implying that these structures are too hot to host microbial life.

In contrast to the active black smoker structures, all other hydrothermal deposits and sediment samples taken meters away from these chimneys (see Supplementary Tables S1, S2 and S4), as well as background sediments ^34^, yielded measurable DNA concentrations (0.14 – 7.2 μg gram^-1^). Porewater geochemical analysis from an AVF sediment core (BC01) shows no consistent hydrothermal signature (Supplementary Table S3). Cl concentrations remain ∼5% below seawater values, but SO_4_ is ∼20% depleted throughout BC01 with no vertical trend. Alkalinity, in contrast, only shows a 20% depletion relative to seawater in the deepest core sections (24-26 cm), while NH_4_ is low, consistently <4µM. It is unclear whether these depletions result from the input of minor quantities of Alk- and SO_4_-poor hydrothermal fluid, or from background processes (e.g. microbial activity, diagenesis) (Supplementary Table S3).

### DNA sequencing and quality filtering

Shotgun sequencing of DNA from all AVF samples positive for DNA resulted in metagenomes with 54.5 – 160M pairs of 150 bp raw sequence reads. Quality filtering passed 89 – 91% of read pairs per sample, resulting in an average of 72.5M (±21M) pairs of high-quality reads per metagenome (Supplementary Table S5). Sequencing of background samples resulted in an average of 31.7M (±3M) high-quality reads with an average of 85 - 88% pairs passing quality filtration per sample.

### Microbial community structure

Taxonomic assignment of metagenome reads mapping to 16S rRNA gene references revealed that all study samples are dominated by Bacteria (77 – 96% of 16S rRNA gene read counts) with comparatively minor representation from Archaea (Fig. 2, Supplementary Table S6). The number of dominant phyla (each comprising ≥1% 16S rRNA gene read abundance of at least one sample) ranges between 8 and 15 bacterial phyla and 0 and 4 archaeal phyla per sample. The most dominant class-level assignment in both active areas and sediment layers above 8 cm depth at AVF was Gammaproteobacteria (17.4 – 50.7%).

**Fig. 2.**
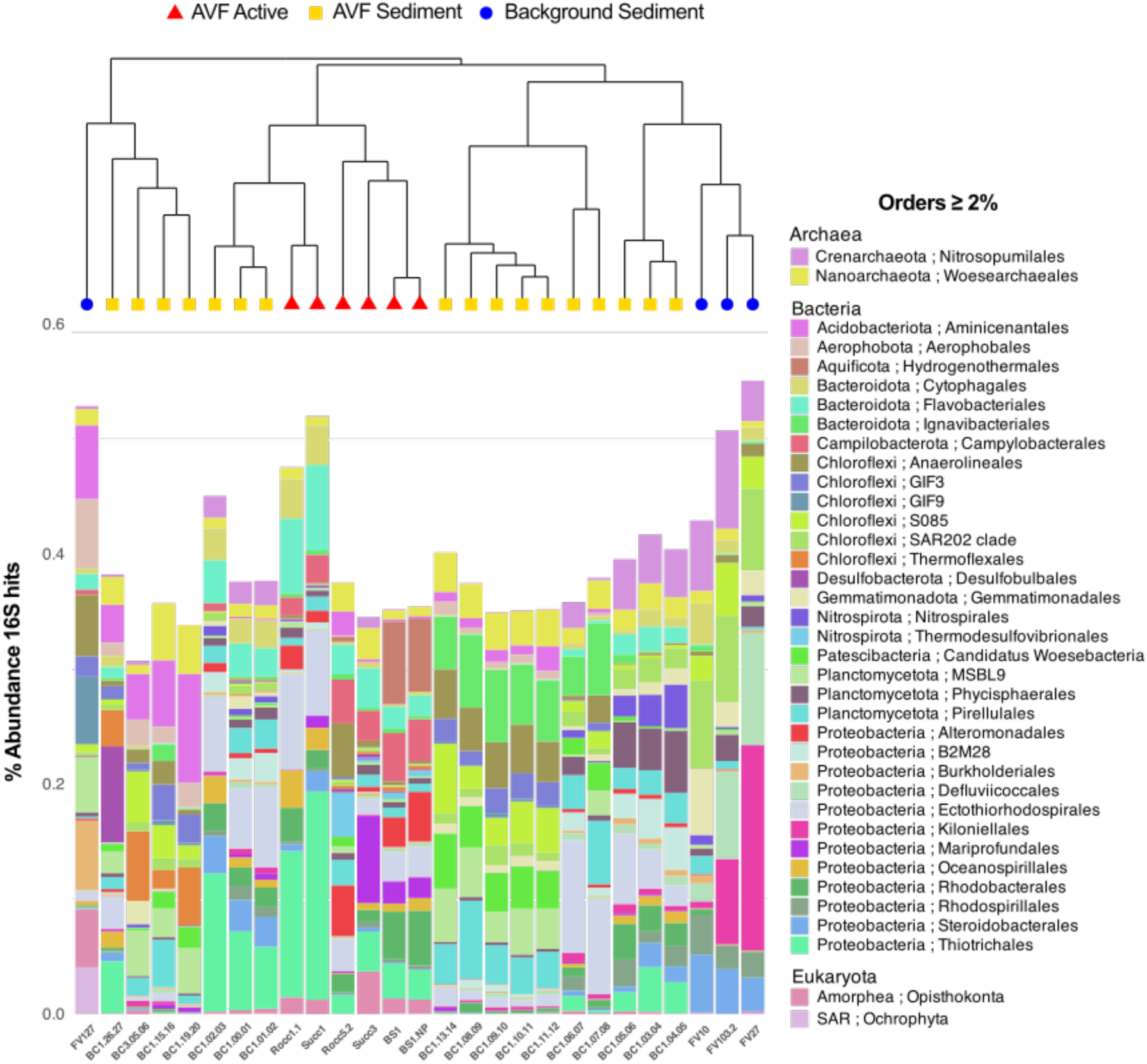
Relative abundance of 16S rRNA gene reads in metagenomes from AVF and background sediments. Samples are grouped by hierarchical clustering based on taxonomic composition. Taxa assigned at order level or lower and representing ≥2% of total 16S reads in at least one metagenome were considered in the analysis.

Bacterial phyla specifically enriched among AVF active samples (Fig. 2) were primarily taxa commonly found at deep sea hydrothermal vents globally ^35^ including Aquificota (7.6%), Campylobacterota (1.5 – 4.2%) and Deinococcota (up to 2.5%). Dominant Pseudomonadota (formerly Proteobacteria) detected specifically at active sites include class Zetaproteobacteria (< 0.1 – 1.9%) and orders Caulobacterales (0.2 – 2.5%; class Alphaproteobacteria) and Methylococcales (< 0.1 – 2.6%; class Gammaproteobacteria). Individual archaeal phyla with relative abundances higher than 1% within active sites include Thermoproteota, Halobacterota, and Nanoarchaeota (see Supplementary Table S9).

To explore whether sediment samples taken meters away from active venting differ from deep sea sediments tens of meters to kilometers away from AVF, we compared the microbial taxonomic distribution between AVF sediment and background sediment samples.

Hierarchical clustering of samples based on order-level community structure (Fig. 2) placed three of the four background samples in a distinct cluster with AVF sediment samples enriched with putative organotrophic bacteria associated with sponges or marine water column and sediment environments, including members of Kiloniellales ^36^, Steroidobacterales ^37,38^, Defluvicoccales ^39,40^, and Chloroflexota orders SAR202 and S085 ^41,42^. Conversely, several dominant taxa only found in AVF surface sediments are associated with marine environments rich in sulfides or heavy metals, including Pseudomonadota orders B2M28 and Ectothiorhodospirales, Rhodobacterales, and Bacteroidota order Flavobacteriales^43–47^.

A total of 1,577 metagenome-assembled genomes (MAGs) with completeness scores greater than 50% and redundancy below 10% with CheckM2 were reconstructed from the metagenomic dataset (Supplementary Table S8). Overall, community structure analyses based on MAG taxonomic assignment and coverage values were highly consistent with 16S rRNA gene analyses (Supplementary Fig. S1 and S2), suggesting that we successfully reconstructed MAGs representing dominating genomes within the most abundant taxa.

### Distribution of genes for autotrophic carbon fixation

The Calvin-Benson Basham (CBB) cycle is the predominant carbon fixation pathway encoded by MAGs recovered from active AVF hydrothermal deposits. This is consistent with the prevalence of Gamma- and Alphaproteobacteria, which are known to use the CBB cycle for CO_2_ fixation ^48^. Campylobacterota and Aquificota MAGs encoding the reverse tricarboxylic acid (rTCA) cycle are limited to AVF hydrothermal deposits (Fig. 2, Supplementary Table S11) while key genes for the Wood-Llungdahl (WL) pathway are found among diverse taxa in all samples. A majority of both Archaea and Bacteria MAGs recovered from AVF samples lack CO_2_ fixation pathways, which suggests that obligate heterotrophs may dominate over facultative or obligate autotrophs in most samples (avg. 65% ± 10.9).

### Key genes for oxidation of reduced sulfur species

We used the presence of Sox cassette genes as evidence for thiosulfate oxidation potential whereas presence of Sox and sulfide:quinone reductase (sqr) or flavocytochrome c sulfide dehydrogenase subunit B (fccB) were used as indicative of H_2_S oxidation potential. Sox genes are most commonly present among abundant Gammaproteobacteria, and to a lesser extent Alphaproteobacteria, at AVF (Supplementary Table S13). Sox-encoding MAGs also include members of Aquificota and Campylobacterota at active AVF sites. Comparatively less abundant and fewer Sox-encoding MAGs occupy background samples (in total 6 MAGs representing .07 - 3.0% of the community) than at AVF sites (in total 168 MAGs representing 0.3 - 67% of the community). A majority of MAGs with Sox system genes have a truncated Sox cassette (*soxYZAB*) rather than the complete set (*soxYZABCD*). Sox genes and sqr are encoded by MAGs that have relative abundances of up to 14% at AVF active sites (Supplementary Table S13), and thus, a majority of the Sox-encoding MAGs at AVF encode the potential to oxidize both sulfide and thiosulfate.

### Iron oxidation

AVF hosts a high diversity of potential iron oxidizing microorganisms, consistent with the typically high dissolved Fe(II) content of ultramafic-influenced hydrothermal fluids analogous to those at AVF ^11,21^. The iron:rusticyanin reductase gene (*cyc2*), an established marker gene for aerobic Fe(II) oxidation, was detected in all high abundance MAGs assigned to Zetaproteobacteria. Members of the bacterial class Zetaproteobacteria (phylum Pseudomonadota) are widespread among iron-rich marine systems ^49–53^ and are comprised primarily of obligate iron oxidizers. Most of the Zetaproteobacteria at AVF encode *cyc2* of clusters 1 or 2, while a few MAGs instead encode *cyc2* of cluster 3. Cluster 3 cyc2 is purported to function in Fe(II) oxidation within Zetaproteobacteria but has a less conserved role among other taxa outside this class ^52,53^. Cyc2 genes of clusters 1 and 2 are also found outside of Zetaproteobacteria at AVF, possibly conferring Fe(II) oxidation ability for some taxa where *cyc2* homologs have been observed previously within Campylobacterota (genus *Sulfurimonas*), and Nitrospirota (class Thermodesulfovibrionia) and individual MAGs within Gammaproteobacteria (genera *Ralstonia*, SZUA-152, SZUA-229) ^53,54^. Interestingly, we also found *cyc2* in MAGs assigned to taxa for which to our knowledge the presence of *cyc2* has not been reported before including Alphaproteobacteria genus *Paremcibacter* (2 MAGs), Nitrospinota class UBA7883 (6 MAGs), and Desulfobacterota class GWC2-55-46 (3 MAGs). While *cyc2* has been observed in other lineages within these broader taxa, its detection in these genera and classes expands the known taxonomic range of putative iron-oxidizing potential.

Consistent with elevated reduced iron availability near venting structures, putative Fe(II)-oxidizing Zetaproteobacteria MAGs and MAGs encoding cluster 1 and cluster 2 cyc2 genes were most prevalent in AVF hydrothermal deposit samples (0.2 – 14% relative abundance) and AVF sediment samples (0.07 – 1.3% relative abundance), whereas putative iron-oxidizers represented a maximum of 0.07% MAG relative abundance in background samples.

### Genes involved in methane consumption and production

Only seven MAGs harboured key genes involved in methane cycling. This includes members of the Gammaproteobacteria genus *Methylohabius* (abundant at the Hans Tore mound sample HC21-ROV16-BS01/BS01-NP), which encode genes for cytochrome c methanol dehydrogenase (mxaF) involved in aerobic methane oxidation ^55^. The only methyl-coenzyme M reductase gene (mcr) identified among all MAGs of this study belongs to an anaerobic methane-oxidizing archaeon (ANME-2) reconstructed from reads obtained from the Hans Tore hydrothermal mound (HC21-ROV16-Succ3).

### Distribution of uptake hydrogenases

The extraordinarily high concentrations of H_2_ in venting fluids at Aurora motivated our investigation of the H_2_ oxidation potential within the microbial community, which we examined through the distribution and abundance of Ni,Fe uptake hydrogenases among reconstructed MAGs.

Ni,Fe uptake hydrogenases of groups 1a – 1f, 1h – 1j and 2a ^56^ were found in a taxonomically wide range of MAGs reconstructed from AVF samples. These detected uptake hydrogenases vary in terms of their H_2_ affinity, sensitivity to oxygen, and what electron acceptors they are associated with ^56^. Both the taxonomic affiliations and functional breadth of uptake hydrogenases indicate that a wide variety of bacteria and archaea potentially use H_2_ as electron donor in aerobic or anaerobic respiration at AVF.

The relative abundance of MAGs encoding one or more uptake hydrogenases in hydrothermal deposit samples taken near vents was consistently above 20 – 30%. Relative abundances of hydrogenase-encoding MAGs were more variable in AVF sediment samples (2.5 - 28%) with the highest abundance seen in the upper sediment horizons (4 - 6 cm depth). All AVF samples had higher abundances of uptake hydrogenase encoding MAGs compared to background samples with the exception of background sample HC19-FV127, where MAGs assigned to phylum Aerophobota encoding group 1a and group 2a hydrogenases were highly abundant (Fig. 3). Group 2a hydrogenases have previously been observed to be involved in either aerobic respiration or recycling of H_2_ from other cellular processes ^56^.

**Fig. 3.**
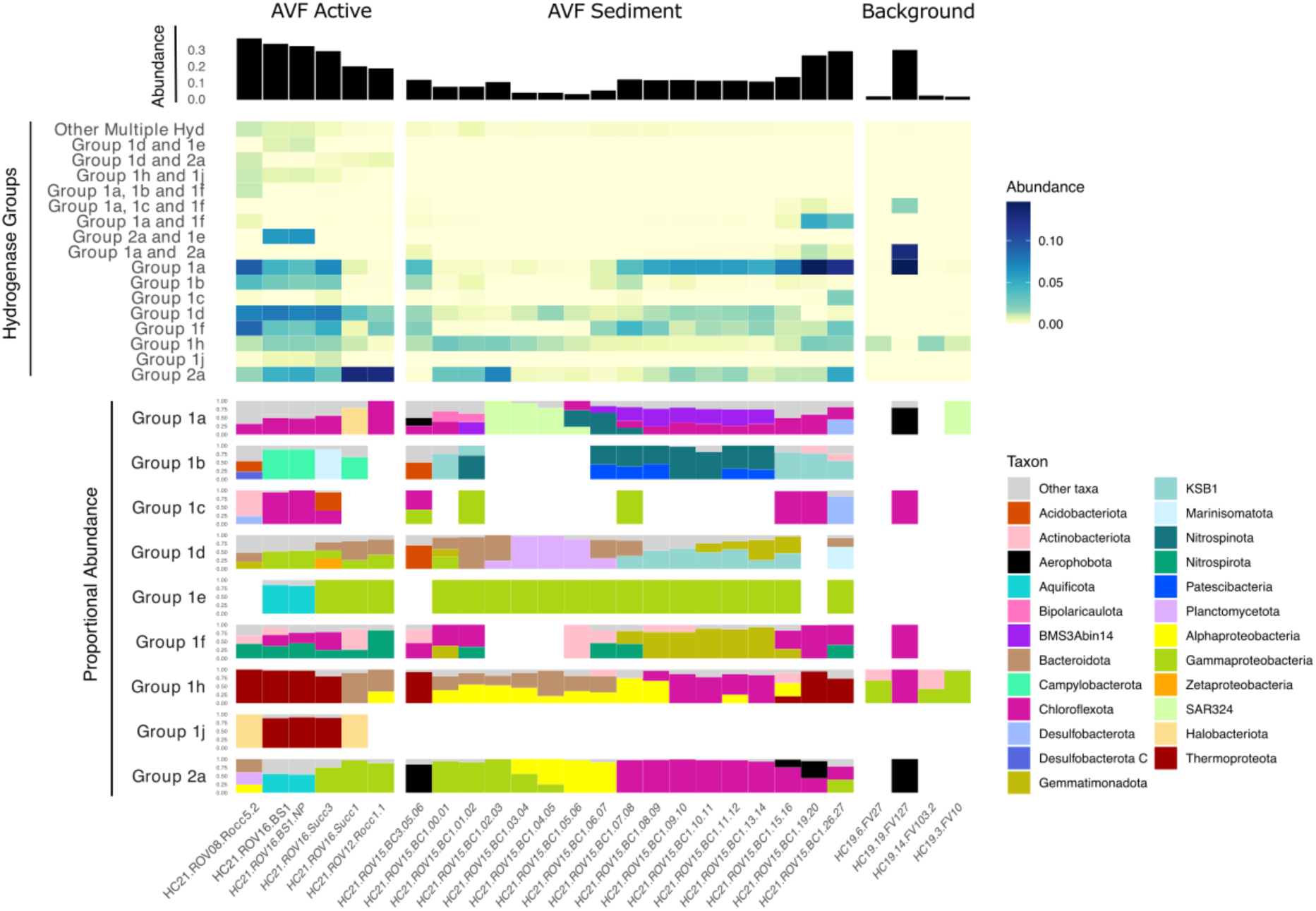
Uptake hydrogenases encoded by MAGs recovered from AVF and background sediment metagenomes. The top panel shows the coverage-based relative abundance of all MAGs with ≥0.5× coverage that encode at least one uptake hydrogenase. The middle panel displays MAG relative abundances partitioned by the hydrogenase group(s) encoded, and the bottom panel depicts the proportional contribution of MAGs from different taxonomic groups to the abundance of each hydrogenase group.

However, upon examination of the genes encoded near the group 2a hydrogenases in the HC19-FV127 MAGs, no hydrogenase maturation related genes are present. In all other background samples only MAGs with hydrogenases of group 1h were detected.

Hydrogenases within group 1h are high-affinity uptake hydrogenases commonly encoded among microorganisms exposed to atmospheric or below-atmospheric levels of H_2_ and are thought to enable microbial cells to use H_2_ oxidation as a supplemental energy source for survival in low-energy systems ^15,57,58^.

Notably, MAGs encoding fast-acting, low affinity hydrogenases (groups 1b and 1d) were only prevalent in active areas and were not detected in background samples (Fig. 3).

Hydrogenases of group 1d are involved in aerobic H_2_ oxidation whereas group 1b hydrogenases are associated with microaerophilic and facultative anaerobic respiration with H_2_ and are tolerant of oxygen ^59^. Group 1b hydrogenases are commonly encoded among facultative aerobes found in hydrothermal systems. At AVF, MAGs encoding group 1b hydrogenases were most abundant in the active sites closest to venting fluids (HC21-ROV16-BS01/BS01-NP, HC21-ROV16-Succ3, HC21-ROV16-Rocc5.2).

Medium and high affinity hydrogenases are also widespread in AVF samples, including hydrogenases involved in aerobic H_2_ oxidation (groups 2a and 1h) and anaerobic H_2_ oxidation connected to reduction of sulfate, metals, and organohalides (group 1f). In most of the background sediment samples (three out of four) all detected uptake hydrogenases belonged to group 1h. Interestingly, more combinations of multiple hydrogenases are encoded by MAGs in active sites than in AVF sediments or in background sediment samples (Fig. 3) indicating a high versatility of H_2_ oxidation modes among organisms inhabiting active sites at AVF.

MAGs encoding uptake hydrogenases at AVF are assigned to a wide range of taxa (Fig. 3). At active sites hydrogenases are widespread in MAGs associated with aerobic autotrophic growth assigned to Gammaproteobacteria, Campylobacterota, Aquificota, and Zetaproteobacteria. Hence, H_2_ oxidation seems to be an important energy source sustaining microbial communities at AVF. We did not, however, detect any obvious example of MAGs derived from obligate H_2_ oxidizers. Rather, lithotrophic primary producers at AVF seem to have capacity for versatile metabolism including the ability to oxidize sulfide or iron in addition to H_2_. MAGs from putative anaerobic sulfate reducers of taxa Nitrospirota, Archaeoglobales, and Desulfobacterota encode oxygen-intolerant uptake hydrogenases typical for these groups. These MAGs are present at active sites, though are only dominant within the deposit sample HC21-Rocc5.2 collected from Enceladus vent, and not at the Hans Tore mound (Fig. 3). While no hydrogenotrophic methanogens were detected at AVF, three genomes of likely H_2_-dependent methylotrophic methanogens of classes Methanomethylicia and Thermoplasmata (order Methanomassilicoccales) were detected at low abundances (Supplementary Table S9). Interestingly, uptake hydrogenases were also widespread among taxa largely associated with organoheterotrophic growth – e.g. Bacteroidota, Chloroflexota, and Planctomycetota.

We describe two MAGs (HC21_M0139 and HC21_M0019) in greater detail below due to their high abundance at AVF active sites, the hydrogenases they encode, and their taxonomic affiliations.

MAG HC21_M0139 was assigned to the genus *Mariprofundus* within class Zetaproteobacteria. *Mariprofundus* is a genus commonly observed in hydrothermal systems around the globe whose members are well known for their ability to oxidize iron ^49^.

Surprisingly, MAG HC21_M0139 (94.4% completeness, 0% contamination) encodes the genomic potential for growing lithotrophically with H_2_ in addition to reduced iron. To our knowledge HC21_M0139 represents the first report of H_2_ oxidation capacity among members of genus *Mariprofundus*. MAG HC21_M0139 encodes all subunits and coenzymes for operational hydrogenases. This includes a group 1d Ni,Fe uptake hydrogenase homolog and a histidine kinase-linked Ni,Fe H_2_-sensing hydrogenase of group 2b (which activates the expression of the respiratory uptake hydrogenase (Supplementary Table S12)). Phylogenetic analyses of the large subunits of these uptake and H_2_-sensing hydrogenases reveal a close evolutionary relationship between *Mariprofundus* hydrogenases and hydrogenases of the Zetaproteobacteria genus *Ghiorsea*, as well as Gammaproteobacteria (Fig. 4, Supplementary Fig. S3 and S4).

**Fig. 4.**
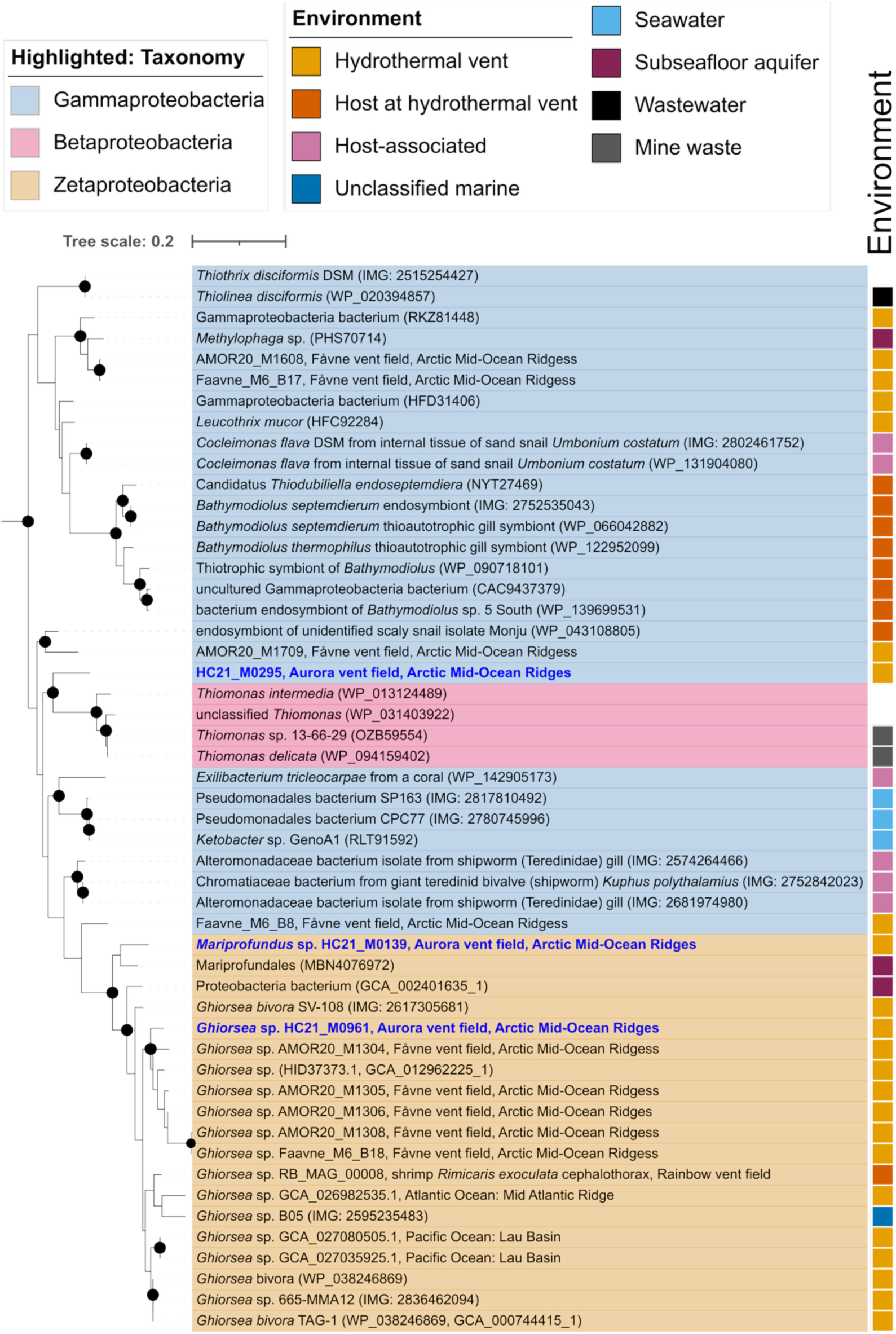
Phylogenetic tree of the large subunit of group 1d Ni,Fe hydrogenases encoded by Zetaproteobacteria MAGs from AVF and all publicly available Zetaproteobacteria genomes, with closest relative reference identified using BLAST. AVF MAGs are highlighted in blue. Black node circles mark branches with support values higher than 80% with SH-like approximate likelihood ratio test and 95% with ultrafast bootstrapping.

MAG HC21_M0019 is assigned to the family *Hydrogenothermaceae* within the phylum Aquificota. Phylogenomic analysis revealed that MAG HC21_M0019 was assigned to placeholder genus JALSSE01 (AAI 73%, Fig. 5, Supplementary Table S10). Two representatives of genus JALSSE01 were previously reconstructed in a global hydrothermal vent metagenomic study ^35^. Surprisingly, the reference MAGs of JALSSE01 do not encode Ni,Fe uptake hydrogenases, despite having 97% and 94% completeness scores with CheckM2 (Supplementary Table S14). MAGs of the neighboring *Hydrogenothermaceae* genus, placeholder JAADDZ01, however, appear to exclusively encode uptake hydrogenases that belong to either aerobic hydrogenase group 2a or Isp-type hydrogenase group 1e. MAG HC21_M0019 encodes both group 2a and group 1e hydrogenases. NCBI BLAST searches of these uptake hydrogenases (accessed November 2025) indicate that both the group 2a and group 1e hydrogenases encoded by HC21_M0019 have closest homology to hydrogenase sequences encoded within MAGs of Aquificota reconstructed from samples taken at deep sea hydrothermal vents. Hydrogenase group 2a has not previously been highlighted as an abundant hydrogenase in hydrothermal systems. Nevertheless, MAG HC21_M0019 is the most dominating MAG at the summit of the Hans Tore mound closest to active venting, representing 7% of the community observed within replicates from the hydraulic cylinder sample (HC21-ROV16-BS01/BS01-NP). Genes encoding all expected subunits and hydrogenase accessory proteins were identified for both hydrogenases encoded by MAG HC21_M0019, including subunit HucE which is critical for H_2_ uptake function in group 2a hydrogenases ^60^. Genomic features-based prediction indicates that HC21_M0019 has an optimal growth temperature of 62.7 °C. This MAG encodes a cbb3-type cytochrome, a terminal electron acceptor for aerobic respiration common to microbes living in low oxygen conditions. The genome also encodes a nitrate reductase, implying that nitrate may serve as an alternative electron acceptor to oxygen. Thus, representatives of MAG HC21_M0019 likely thrive close to active venting with high concentrations of H_2_ and possess hydrogenases consistent with both aerobic and anaerobic respiration, suggesting a flexible hydrogenotrophic lifestyle.

**Fig. 5.**
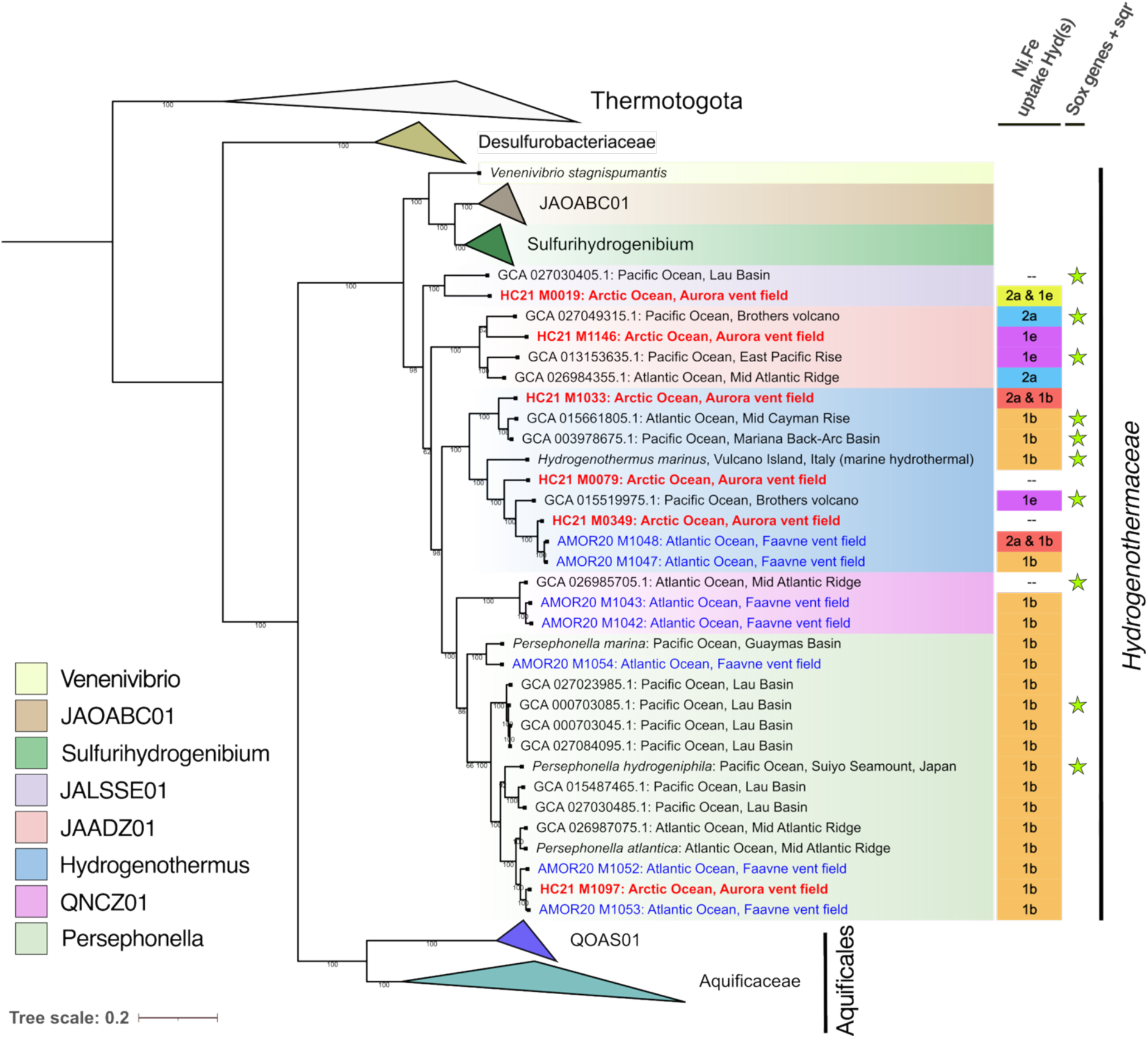
Maximum-likelihood phylogenomic tree of MAGs from this study and publicly available genomes assigned to the Aquificota, with *Thermotoga* as the outgroup. The tree is expanded to highlight the family *Hydrogenothermaceae*, to which all AVF MAGs were assigned. AVF MAGs are highlighted in red, and Fåvne vent field MAGs in blue. Encoded uptake hydrogenases are labeled, and the presence of *sox* and *sqr* genes is indicated by green stars.

## Discussion

Ultramafic-influenced hydrothermal vents are rare within the current inventory of known vent fields ^61^ and remain sparsely characterized microbiologically ^31,62–70^. The Aurora Vent Field thus adds an important new ecosystem to this limited knowledge base. Using genome-resolved metagenomics, we characterized the microbial community of AVF, revealing functional properties of distinct microbial lineages adapted to this H_2_-rich setting. Source fluids at AVF are exceptionally enriched in H_2_ ^25,26^, potentially creating one of the most energy-rich deep-sea habitats yet described. Hence, AVF represents a unique natural laboratory to investigate how extremely high H_2_ availability shapes microbial communities and chemosynthetically driven food webs.

Although metagenomic studies in deep-sea hydrothermal systems are becoming more common, most previous work has relied on 16S rRNA gene surveys. Despite the detailed information gained from these studies on the distribution of taxonomic groups, the metabolic potential of many of the microorganisms found in hydrothermal systems, and the flow of carbon and energy through microbial food webs, remains poorly characterized. H_2_ availability has been shown to influence the diversity of hydrogenases in hydrothermal systems ^71^, yet the breadth of uptake hydrogenases at vents is rarely assessed. Previous reports often focus on particular chemoautotrophic community members ^72–75^, infer H_2_-oxidizing potential from taxonomic assignment ^76^, or document only low-affinity uptake hydrogenases ^35,68,77,78^.

Through reporting of all predicted uptake hydrogenases in this work, we assess the total H_2_ consumption potential at AVF. As discussed below, our results provide several clues about how H_2_ may contribute to shaping microbial communities at AVF.

### Hydrogen oxidizers at AVF encode versatile respiration potential

Interestingly, despite the high energetic potential for H_2_ oxidation at AVF, we see no evidence for the presence of obligate H_2_ oxidizers in any sample. Rather, H_2_ oxidation seems to be a supplemental energy source for a wide range of organisms. Hence, versatility in substrates used for energy acquisition seems to be favoured over obligate H_2_ oxidation even under extremely H_2_-rich conditions. Arguably, microorganisms living in mineral deposits and biofilms in hydrothermal systems could experience frequent and rapid shifts in substrate availability due to variations in fluid flow. At Aurora, even minute fractions of wafting endmember fluid could contain significant H_2_; for a 10^4^ diluted endmember, H_2_ could still be present at levels of a few micromolar if mixed conservatively. Ecological theory suggests that such variations favour metabolically flexible organisms, where generalists are favoured in environments with high substrate variability over space and time ^79,80^. In this context it is also intriguing that MAGs obtained from AVF samples overall encode a wider range of uptake hydrogenases and a higher number of combinations of these hydrogenases than MAGs obtained from background sediments (Fig. 3). This may also reflect a high versatility in how H_2_ is consumed at AVF and that this versatility occurs as a response to high H_2_ availability (albeit at variable levels).

### Aurora is a hydrogenase hotspot

Compared to background sediments, AVF samples host microbial communities with an overall higher abundance of hydrogenases, a higher diversity of hydrogenases, and a considerably higher diversity of taxa encoding hydrogenases (Fig. 3). Consistent with previous findings from hydrothermal systems ^17,81–83^, we found low-affinity uptake hydrogenases were exclusively encoded by MAGs from active sites. Based on this, AVF can be considered a potential hotspot for hydrogenase diversity and H_2_ oxidizers both in terms of abundance and taxonomic breadth.

The presence of a hydrogenase in iron-oxidizing *Mariprofundus* (Zetaproteobacteria) is intriguing. *Mariprofundus* have been sampled repeatedly at hydrothermal vents, however this is the first time that uptake hydrogenase has been observed within genomes belonging to this genus. The H_2_-rich environment at AVF may select for strains with H_2_ oxidation capability that otherwise exist as low-abundance lineages that typically remain undetected in lower-H₂ settings. Alternatively, the hydrogenases may have been acquired through horizontal gene transfer. In the latter case, the close sequence similarity of the identified hydrogenases to homologs within the broader phylogenetic group (other Zetaproteobacteria) supports transfer among closely related lineages. That this lineage has thus far only revealed H_2_ oxidation potential in an extremely H_2_-rich vent field raises the possibility that the unprecedented H₂ availability plays a role in the emergence or selection of H_2_-oxidizing variants within *Mariprofundus*.

Given the abundant H_2_ in the fluids, it is surprising that we find no evidence for the presence of hydrogenotrophic methanogens in any sample. Abundances of potentially hydrogenotrophic sulfate reducers are also low. Possible reasons for this are that methanogens and sulfate reducers are strictly anaerobic and easily inhibited by the influence of oxic seawater. Methanogens are widely present in numerous other hydrothermal systems, particularly in H_2_-rich systems, and even within diffusely venting areas ^84^, and are typically found in the well-developed interior gradients of chimney structures. It seems likely that methanogens are also present at AVF, but that our samples do not include strictly anaerobic zones where methanogens can thrive.

### Hydrogenothermaceae at AVF may be adapted to dynamic H2-rich conditions

It is interesting to note that abundant members of the Aquificota phylum present in AVF are highly related to placeholder genera within the *Hydrogenothermaceae* family with several members which appear to encode group 2a hydrogenases as their primary uptake hydrogenases (Fig. 4). Group 2a hydrogenases are typically characterized as high-affinity aerobic hydrogenases that enable growth under trace-H₂ conditions, generally co-occurring with low-affinity group 1b/d hydrogenases in terrestrial Aquificota ^58^. A functional study further showed that the group 2a hydrogenase of the model bacterium *Mycobacterium smegmatis* is constitutively expressed across a wide range of H_2_ partial pressures ^85^. This is in opposition to low-affinity groups 1b/d or high-affinity group 1h hydrogenases, which become less active when H_2_ partial pressure is outside a certain range ^86–88^. Hydrogenases of group 2a are also reported to have a high catalytic capacity and the ability to oxidize H₂ efficiently across a wide range of concentrations ^89,90^. However, group 2a hydrogenases are not considered as primary hydrogenases in high-temperature, ultramafic influenced deep-sea hydrothermal systems, where H_2_ is abundant but may rapidly fluctuate. Their presence as the dominant uptake hydrogenases in *Hydrogenothermaceae* at AVF suggests that they may support H_2_ respiration across dynamic spatial and temporal gradients in H₂ availability in vent settings, expanding the known ecological role of group 2a hydrogenases.

### EHect of hydrogen oxidation on the flow of carbon and energy at AVF

It is well known that numerous heterotrophic organisms have the ability to acquire energy from H_2_ oxidation ^91^. This includes obligate organoheterotrophs that use H_2_ oxidation as a supplementary energy source. The distribution of H_2_-utilizing organotrophs in hydrothermal systems, however, is not well documented. Here, we show that uptake hydrogenases are widespread in putative organotrophic organisms at AVF, including highly abundant MAGs of Bacteroidota, Planctomycetota, and Chloroflexota, phyla in which hydrogenases have been previously detected among some members collected from various environments including deep sea hydrothermal sites ^56,58,92–95^. If many of the organotrophs at AVF use H_2_ oxidation as a supplemental energy source in mixotrophy, this would allow them to use more of the carbon from primary production for biomass rather than energy acquisition, hence making the flow of carbon from primary producers to consumers more efficient ^94^. This might partly explain how a relatively small community of primary producers, estimated to be 18-55% of the community in active sites at AVF (based on MAGs encoding key genes for CO_2_ fixation, Supplementary Table S10), can sustain a larger community of heterotrophic consumers.

### AVF sediments

It is difficult to assess how the H_2_-rich fluids may influence sediment communities surrounding mounds and chimneys at AVF. On the one hand, influence from diffuse hydrothermal fluids is not evident from the porewater geochemistry data. However, the relatively high diversity and abundance of uptake hydrogenases in the upper centimeters of AVF sediments may point to intermittent exposure to vent-derived H_2_ (Fig. 3). We cannot preclude minute contributions of H_2_-rich vent fluid being present that are not detectable (akin to the CH_4_ ‘ground fog’ or ultra-diffuse flow inferred in other sites of serpentinization ^96^). Alternatively, the communities here may be solely supported by sedimentation of organic-rich material, including biomass produced in hydrothermally influenced areas such as the plume. In the latter case, the uptake hydrogenases present in the sediments may be used mainly to exploit H_2_ produced via fermentation. A parallel investigation into these same sediment cores indicated that there is a peak in organic matter in the top 7 cm of the cores, with signatures of lithotrophy ^34^. Thus, AVF sediment communities may, at a minimum, be indirectly connected to high H_2_ concentrations in venting fluids through enhanced productivity at active sites.

### Community organization under extreme H2 availability

Our metagenomic analyses suggest that the AVF communities are largely driven by H_2_ oxidation and sulfide oxidation. The high H_2_ hydrothermal fluids at AVF appear to sustain a community characterized by diverse, abundant taxa capable of H_2_ oxidation using a varied suite of uptake hydrogenases. Supplementary H_2_ oxidation in organotrophic organisms at AVF may influence the flow of carbon and energy in the microbial communities, potentially allowing a relatively low number of autotrophic primary producers to sustain a larger heterotrophic biomass. To further assess how H_2_ availability shapes communities and genetic features of microorganisms present in AVF, and in deep sea hydrothermal systems more broadly, more knowledge is needed about the overall diversity and distribution of hydrogenases, the transcription of and expression profiles of hydrogenases, *in situ* H_2_ oxidation rates, and the H_2_ oxidation potential of individual strains.

## Materials and Methods

### Sampling

Seafloor sampling was carried out by the Norwegian research icebreaker RV Kronprins Haakon in October, 2021 at AVF (82°53’49’’N, 6°15’21’’W), using the work-class ROV *Aurora* provided by REV Ocean as described in detail in Ramirez-Llodra et al. (2023)^23^. Materials were recovered using either a spring-loaded box corer (aka. blade corer) for sediments, a vacuum suction tool for loose hydrothermal deposit material, a hydraulic cylinder for biofilms, or simply the manipulator itself for fragments of hydrothermal deposit (Supplementary Table S1). Sediment blade core horizons (<30 cm deep) were subsampled throughout in a 4 °C cold room using ethanol-cleaned steel tools. Sampling of sediments with a multicorer at locations distal to AVF carried out in 2019 on the same vessel were processed similarly (Supplementary Table S2). Ten hydrothermal deposit fragments were collected from chimney structures and subsampled for microbiology using a sterile scalpel by scraping ∼1 cm deep, from outer to inner chimney wall sections. All microbial samples were stored in cryotubes or sterile sampling bags, and immediately flash frozen to -80 °C onboard the vessel until further processing.

### Porewater Analysis

Porewater samples were obtained from AVF sediment core BC01 (28cm, at 2cm intervals), located ∼4.5m from the Enceladus vent ^34^, using Rhizon samplers ^97^. The first 2-3 mL of extracted porewater was discarded to minimize potential contamination with rinsing ultrapure water. Samples were processed under cold conditions, transferred to Nalgene HPDE vials, and stored appropriately. Alkalinity (Gran titration) and pH (combination electrode) measurements were performed on board after sampling porewaters. Aliquots for anion and ΣNH_4_ analyses were stored frozen until laboratory analysis by high pressure ion chromatography, as described in Pereira et al. (2025) ^98^.

### DNA Extraction and Sequencing

The DNeasy Powersoil kit (Qiagen, Germany) was used to extract DNA according to the manufacturers specifications, utilizing ∼0.5 g of soil or chimney sample as input for extraction. In the case where sample DNA concentrations were low, the Powersoil Max DNA extraction kit used to recover a higher DNA yield from up to 10 g of sample. All extractions were stored in lo TE Buffer (10 mM Tris, 0.1 mM EDTA) and sent for sequencing. Shotgun metagenomic libraries from AVF samples were sequenced using Illumina NovaSeq (150 bp paired-end) at the Norwegian Sequencing Centre (Oslo, Norway). Background sediment samples were sequenced using Illumina HiSeq (150 bp paired-end).

### Sequence data processing

Demultiplexed metagenomic Illumina reads were processed following the procedures described in Hribovšek et al. (2023) ^31^, with minor modifications. Briefly, raw sequence quality was assessed using FastQC v0.12. ^99^ and MultiQC v1.13 ^100^. Potential human DNA contamination was evaluated by mapping reads against a human genome reference mask (hg19) using BBMap ^101^. Adapters were trimmed with Trimmomatic v0.38 ^102^ and read quality trimming and filtering were performed using wrapper scripts implemented in Anvi’o v7^103^.

To assess community composition, quality-controlled reads from metagenomes were mapped entries in the SILVA 16S rRNA reference database ^104^ using PhyloFlash v3.0 ^105^, where read best-hits determined taxonomic assignment based on match identity.

Assembly was conducted using MEGAHIT v1.2.9 ^106^, generating contigs ≥1,500 bp. Single-sample read mapping to assembled contigs using Bowtie 2 v2.4.2 ^107^ was followed by genome binning implemented in Anvi’o using MetaBAT 2 v2.12.1 ^108^, MaxBin 2 v2.2.4 ^109^, and CONCOCT v1.1.0 ^110^; consensus bins generated were selected using DASTool v1.1.4 ^111^. Bins created from each metagenome were dereplicated to 98% identity using dRep v3.2.0 ^112^ to create a non-redundant set of bins across samples. Bin completeness and redundancy were estimated with CheckM 2 v1.1.0 ^113^, and bins with >50% completeness and <10% redundancy were retained as metagenome-assembled genomes (MAGs) for downstream analyses.

Quality-controlled reads from each sample were subsequently mapped to the non-redundant MAG dataset using Bowtie2 to estimate relative MAG abundances based on genome coverage. Taxonomic classification of MAGs was assigned using GTDB-Tk v2.3.2 ^114^.

Pairwise genomic relatedness was assessed using pyANI v0.2.12 ^115^ (implemented in Anvi’o with *anvi-compute-genome-similarity*) and ezAAI v1.2.2 ^116^. Species and genus boundaries were defined at ∼95 % ANI and 65 % AAI ^117,118^.

Gene prediction and annotation of MAGs were performed using a modified version of the pipeline originally described by Dombrowski et al. (2020) ^119^. Briefly, gene calling and preliminary functional annotation were carried out with Prokka v.1.14.6 ^120^, and subsequent HMM searches were conducted against the COG ^121^, arCOG ^122^, KEGG ^123^, SignalP ^124^, and CAZy ^125^ databases. Hydrogenases were verified through HydDB ^56^, CO₂ fixation genes through KEGG Decoder v1.2.1 ^126,127^, iron and manganese oxidation genes via FeGenie v1.1 and MagicLamp v1.0 ^128,129^, and metal-resistance genes using the BacMet v2.0 database ^130^. Predicted optimal growth temperatures were inferred from genomic features using regression models trained on bacterial genomes ^131^.

### Phylogenomic Analyses

#### Zetaproteobacteria

To place AVF Zetaproteobacteria in a broader phylogenomic context, all publicly available genomes assigned to this class (n = 73; taxid 580370) were downloaded from NCBI GenBank using ncbi-genome-download v0.3.0 (September 2023). Additional genomes were obtained from the Earth’s Microbiome Project ^132^, JGI IMG, and previously published datasets ^53,133–135^. Campylobacterota and Alphaproteobacteria were included as outgroups. Only genomes of at least medium quality (> 50 % completeness, < 10 % contamination) were retained.

Twelve manually curated single-copy core genes for domain Bacteria ^31,103^ were aligned with MAFFT L-INS-I v7.397. The resulting alignments were concatenated then trimmed with trimAl v1.4. rev15 ^136^ with the -gt 0.5 -cons 60 trimming option. Maximum-likelihood phylogenomic trees were inferred with IQ-TREE v2.0.3 ^137^ using the LG+F+R9 substitution model selected in ModelFinder ^138^, with 1,000 SH-like approximate likelihood ratio test replicates ^139^ and 1,000 ultrafast bootstrap replicates ^140^. All phylogenies constructed in this study were visualized and annotated using iTOL v6 ^141^.

#### Aquificota

The phylogenomic tree for Aquificota was constructed to place AVF Aquificota MAGs within known Aquificota relationships, using Thermotogota as the outgroup. Reference genomes were retrieved from NCBI Genbank and JGI databases, dereplicated at 100% identity with dRep, and evaluated with CheckM2; only genomes with > 80% completeness and < 5% redundancy were retained. Taxonomy was assigned using GTDB-Tk, and reference genomes failing the tools quality thresholds were removed. For each species-level group, the genome with the highest completeness and lowest redundancy was selected as the representative. The phylogenomic tree was built using a custom pipeline wrapper script that utilizes VBCG, a set of 20 validated bacterial core genes for producing phylogenies with high phylogenetic fidelity and resolution ^142^. Briefly, VBCGs were identified using hmmsearch on protein sequences predicted by Prodigal, aligned with MAFFT, and trimmed with trimAl. Maximum-likelihood trees were inferred in RAxML v8.2.12 ^143^ under the LG substitution model with CAT rate heterogeneity and automatic bootstrap convergence. Genomes missing > 50% of queried HMMs and marker genes present in < 50% of genomes were excluded. Pipeline scripts are available on GitHub: https://github.com/hakonda2/Phylogeny_PPCM/tree/main.

### Single-gene phylogenies

For NiFe hydrogenases, large-subunit sequences of groups 1d (hya) and 2b (hup) were retrieved from AVF MAGs, reference Zetaproteobacteria, and close relatives identified by NCBI protein BLAST (October 2023). Sequences were aligned with MAFFT, curated in AliView v1.26 ^144^, trimmed with trimAl, and analyzed in IQ-TREE under best-fit LG models. Alignments shorter than 460 aa (hya) or 400 aa (hup) were excluded. Trees were midpoint-rooted, and environmental metadata were sourced from NCBI and GTDB.

## Acknowledgements

The authors would like to acknowledge the captain, officers, and crew of the RV Kronprins Haakon, and the REV Ocean ROV Aurora team, for their pioneering efforts to facilitate seafloor sample collection in challenging ice conditions. We thank Eva Ramirez-Llodra and Stefan Buenz for organizing and leading the 2019 and 2021 HACON research cruises. We would also like to thank the R and the Anvi’o development teams for the open-source packages and documentation they publicly provide that made this analysis and visualization of metagenomic data repeatable and accessible. Funding for onboard sample collection in 2019 and 2021 and science team meetings was supported under the FRINATEK HACON project funded by the Research Council of Norway (project no. 274330). Extraction and sequencing funding was provided by the DeepSequence Project (project no. 315427) as well as a Meltzer Foundation PhD project grant awarded to E. Denny in 2021 (project no. 102737106).

## Author Contributions

**Emily Olesin Denny** – study conception (lead), original manuscript draft (lead), analysis (lead), sampling (lead). **Petra Hribovšek**– original manuscript draft (supporting), phylogenetic analysis (lead), manuscript review (supporting). **Samuel I. Periera** –sampling (supporting), analysis (supporting), manuscript review (supporting). **Claudio Argentino** – analysis (supporting), manuscript review (supporting). **Guiliana Panieri** – sampling (lead), study conception (supporting), manuscript review (supporting). **Achim Mall –** sampling (supporting), analysis (supporting), manuscript review (supporting). **Francesca Vulcano** – sampling (supporting), analysis (supporting), manuscript review (supporting). **Eoghan P. Reeves** – sampling (supporting), project administration (supporting), analysis (supporting), manuscript review (supporting). **Runar Stokke** – project administration (lead), original draft (supporting), manuscript review (supporting). **Ida H. Steen** – sampling (lead), study conception (supporting), project administration (supporting), original manuscript draft (supporting), manuscript review (supporting). **Håkon Dahle** – study conception (lead), project administration (lead), original manuscript draft (supporting), manuscript review (supporting).

## Data Availability

The MAG datasets generated and analysed during the current study are available at NCBI under BioProjects PRJNA1363180 and PRJNA949439. See Supplementary Table S7 for individual MAG BioSample accession numbers.

