## Supplemental Information for "Aurora vent field is a hotspot for microbial hydrogen oxidation in the Arctic Ocean"

### Supplementary Tables

Supplementary Tables S1–S5 are provided in Supplementary_Data_1 (Excel format).
Supplementary Tables S6–S14 are provided in Supplementary_Data_2 (Excel format).

***Supplementary Data 1 (Excel):***
Table S1 – Metadata for seafloor microbial sampling on expedition HACON21
Table S2 – Metadata for microbial sampling on expedition HACON19
Table S3 – Porewater geochemical analyses from sediment core BC01 near Enceladus vent
Table S4 –DNA extraction results from HACON21 and HACON19 microbial samples
Table S5 – Read statistics for Illumina NovaSeq/ HiSeq sequencing

***Supplementary Data 2 (Excel):***
Table S6 – Metagenome sample reads mapping to 16S / 18S rRNA sequences i
Table S7 – Taxonomic assignment of study MAGs
Table S8 – study MAG completeness and redundancy estimates
Table S9 – Relative abundances of MAGs among samples
Table S10 – *Hydrogenothermaceae* average amino acid identities (AAI)
Table S11 – Key and supporting genes involved in microbial $CO_2$ fixation pathways
Table S12 – sequence hits to hydrogenase HMMs in HydDB
Table S13 – Key and supporting genes in study MAGs for lithotrophic processes
Table S14 – Metadata for reference genomes used in the Aquificota phylogenomic tree

### Supplementary Figures

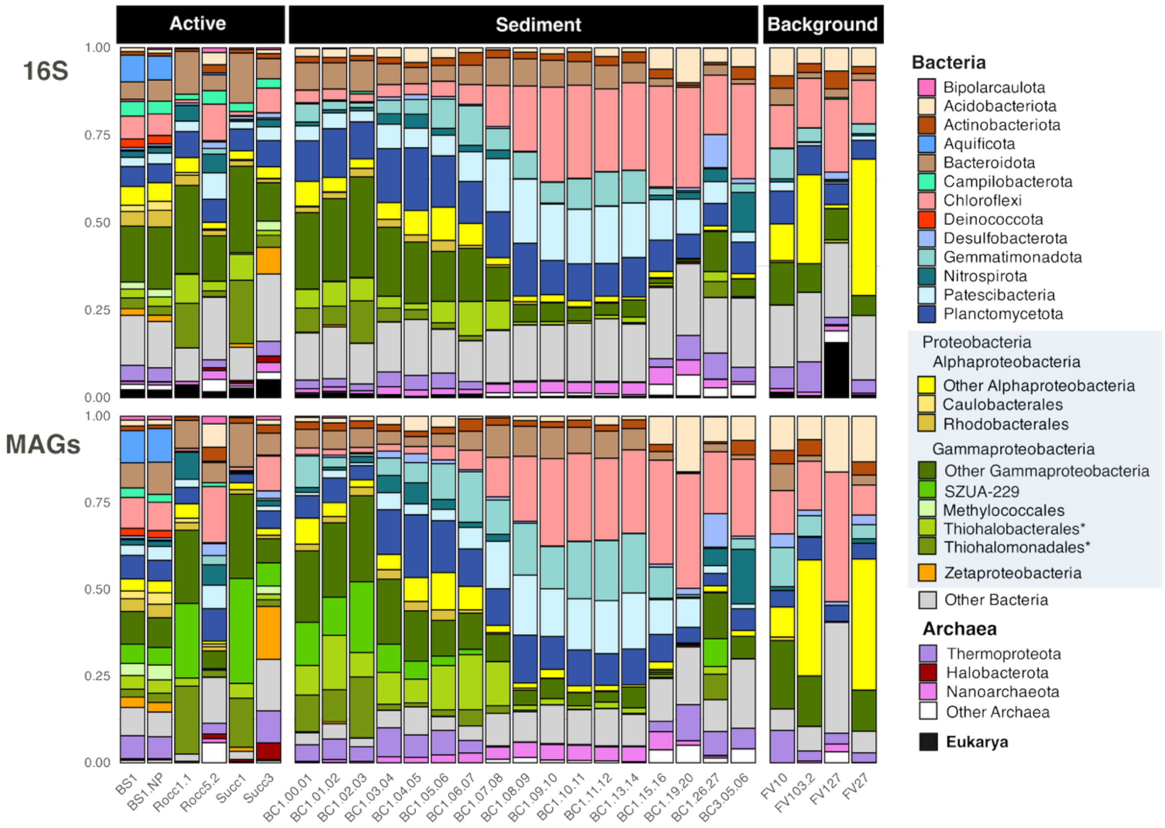


**Figure S1.** Stacked bar plots showing the relative abundances of microbial taxa based on reads mapping to 16S rRNA genes (top panel, input data from Supplementary Table S6) and on MAG coverage per sample (bottom panel, input data from Supplementary Tables S7,S9). In the bottom panel, relative abundances represent the proportion of total MAG coverage attributed to each taxon within a sample. Starred groups Thiohalobacterales and Thiohalomonadales were assigned to orders Alteromonodales and Ectothiorhodospirales respectively based on 16S sequences alone.

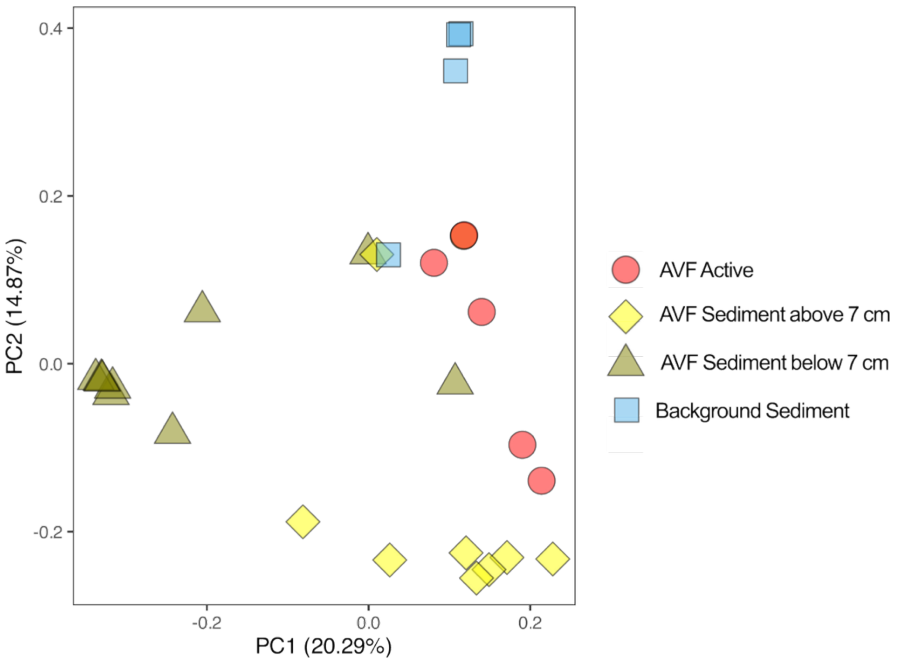


**Figure S2. Principal component analysis of AVF and background samples based on the relative abundance of MAGs using coverage (related to Supplementary Table S9).** Groups of horizons of AVF sediments are separated by 7 cm depth cutoff to highlight an apparent shift in the community diversity at this depth.

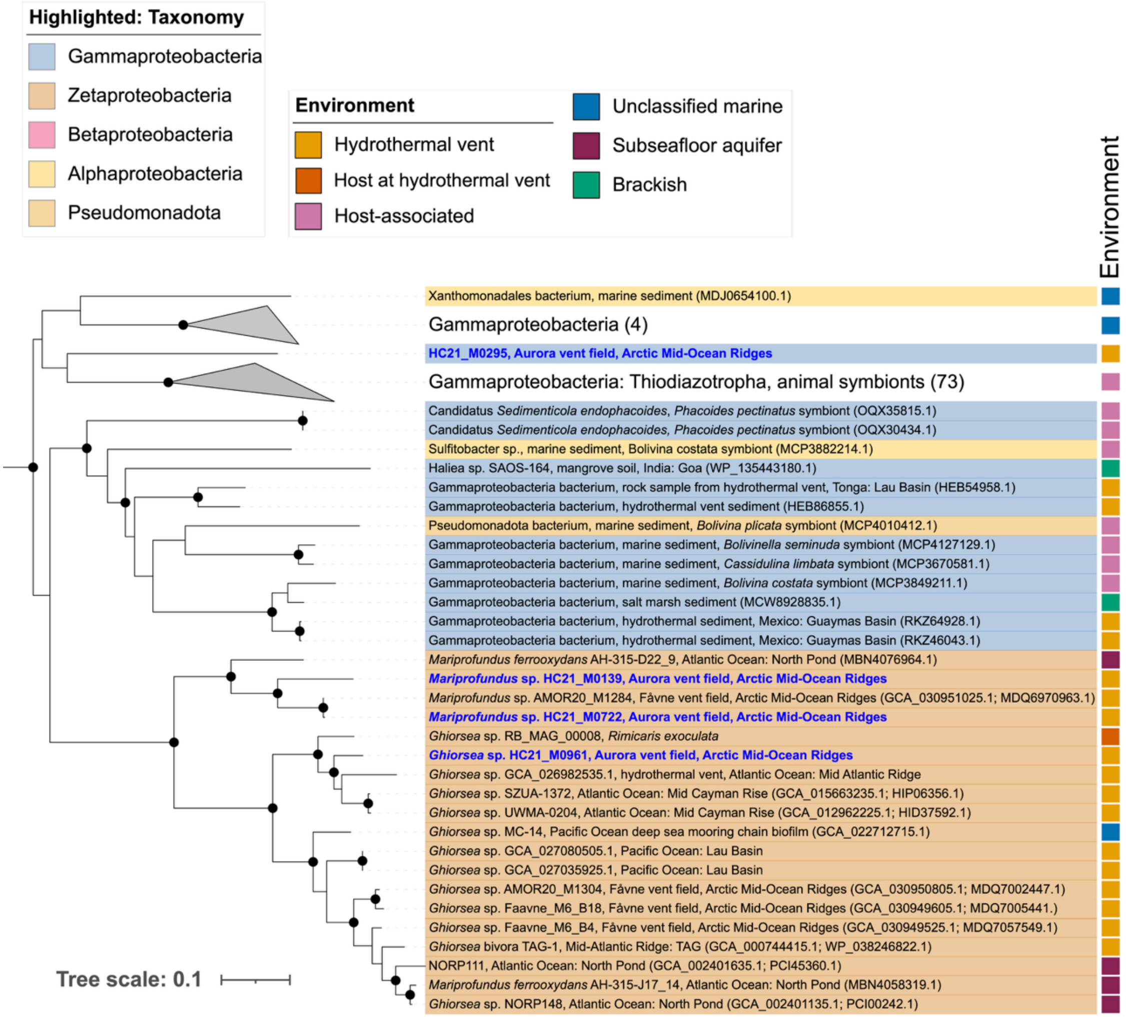


**Figure S3. Phylogenetic tree of the large subunit of group 1d Ni,Fe hydrogenases encoded by Zetaproteobacteria.** The tree includes hydrogenases encoded by Zetaproteobacteria MAGs from AVF, hydrogenases identified in all publicly available Zetaproteobacteria genomes, as well as closest relative references using BLAST. AVF MAGs are highlighted in blue. Black node circles mark branches with support values higher than 80% with SH-like approximate likelihood ratio test and 95% with ultrafast bootstrapping.

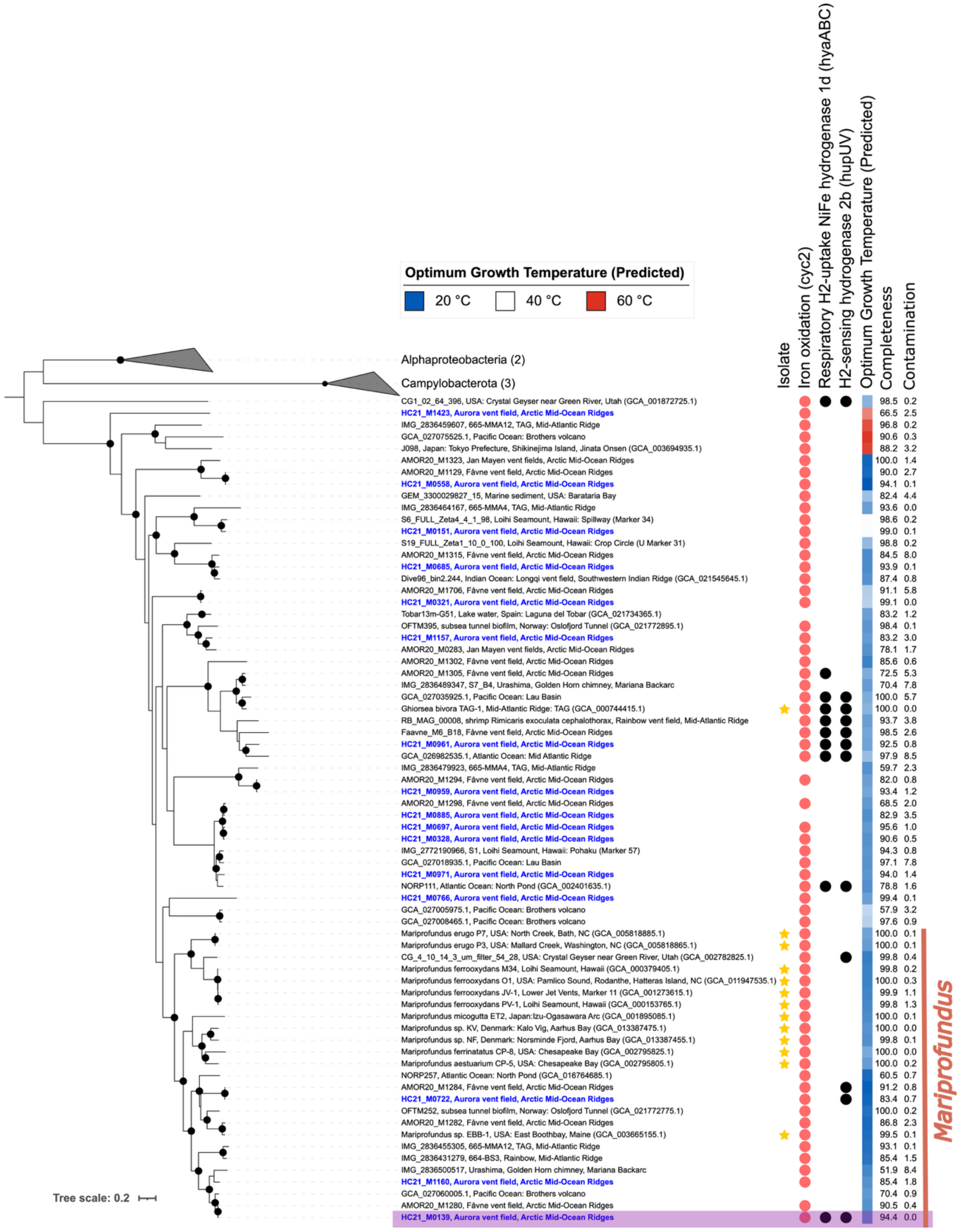


**Figure S4. Maximum likelihood phylogenomic tree of Zetaproteobacteria from AVF**. The tree is based on a concatenated alignment of a manually curated set of 12 single copy gene markers using MAGs from this study and references. AVF MAGs are highlighted in blue. Black node circles mark branches with support values higher than 80% with SH-like approximate likelihood ratio test and 95% with ultrafast bootstrapping.
